# Meta-analysis of massive parallel reporter assays enables functional regulatory elements prediction

**DOI:** 10.1101/202002

**Authors:** Anat Kreimer, Zhongxia Yan, Nadav Ahituv, Nir Yosef

## Abstract

Deciphering the potential of non-coding loci to influence gene regulation has been the subject of intense research, with important implications in understanding genetic underpinnings of human diseases. Massively parallel reporter assays (MPRAs) can measure regulatory activity of thousands of DNA-sequences and their variants in a single experiment. With increasing number of publically available MPRA datasets, one can now develop data-driven models which, given a DNA-sequence, predict its regulatory activity. Here, we performed a comprehensive meta-analysis of several MPRA datasets in a variety of cellular contexts. We first applied an ensemble of methods to predict MPRA output in each context and observed that the most predictive features are consistent across datasets. We then demonstrate that predictive models trained in one cellular context can be used to predict MPRA output in another, with loss of accuracy attributed to cell-type specific features. Finally, we show that our approach achieves top performance in the Fifth Critical Assessment of Genome Interpretation “Regulation Saturation” Challenge for predicting effects of single nucleotide variants. Overall, our analysis provides insights into how MPRA data can be leveraged to highlight functional regulatory regions throughout the genome and can guide effective design of future experiments by better prioritizing regions of interest.

## INTRODUCTION

Massively Parallel Reporter Assays (MPRA) (Weingarten-Gabbay and Segal, 2014) provide cost effective, high-throughput activity screening of thousands of sequences and their variants for regulatory activity (Kheradpour, et al., 2013; Melnikov, et al., 2012; Mogno, et al., 2013; Patwardhan, et al., 2012; Sharon, et al., 2012; Smith, et al., 2013). In these assays, a library of putative regulatory elements is cloned and then transfected or infected into cells of interest. Each element is either associated with a unique barcode or can serve as a unique barcode itself (Arnold, et al., 2013). The activity associated with each given regulatory element (i.e. MPRA output) is assessed by sequencing the transcribed barcodes and estimating the ratio between the transcribed RNA and the construct’s DNA. Since MPRA is still a nascent technology, the development of computational tools that take advantage of existing MPRA datasets could help improve future MPRA candidate sequence selection, enhance our ability to predict functional regulatory sequences, and increase our understanding of the regulatory code and how its alteration can lead to phenotypic consequences.

Previous work have used single MPRA datasets to better identify functional DNA sequences and then study the features that make a sequence regulatory active (Grossman, et al., 2017; Lee, et al., 2015; Sharon, et al., 2012). For example, in the expression quantitative trait loci (eQTL) causal SNP challenge of the Fourth Critical Assessment of Genome Interpretation (CAGI4) community experiment, participants developed methods for predicting regulatory activity of candidate genomic regions and the effect of minor variants on their regulatory potential in MPRA (Beer, 2017; Kreimer, et al., 2017; Zeng, et al., 2017). The main lessons learned from this community effort highlighted the effectiveness of ensembles of non-linear methods, especially when used on features related to transcription factor (TF) binding and chromatin accessibility. Interestingly, epigenetic properties predicted from DNA sequence (Alipanahi, et al., 2015; Zeng, et al., 2016; Zhou and Troyanskaya, 2015) were shown to be more predictive features than experimentally measured epigenetic properties.

While these efforts provided an important first step, each of them focused on a single MPRA dataset in a specific cellular context. Critical questions therefore remain as to how generalizable the insights from MPRA experiments are to other datasets or other cellular contexts. Here, we present a first comprehensive analysis of several MPRA datasets collected by different labs and in various cellular systems; these datasets explore the effect of endogenous loci in several different cell types. We derive a large set of properties to characterize each putative regulatory region and compare the performance of different methods and features for predicting MPRA output. We show that MPRA activity is predictable and that prediction methods tend to perform consistently well when tested on different datasets, with better performance for non-linear methods and favorable results when using an ensemble approach. Consistently, the predictive capacity of individual features is comparable across datasets, with transcription factor binding and epigenetic properties being the top predictors.

We next turned to investigate the generalizability of our models across datasets, which allowed us to distinguish between determinants of MPRA activity that are dependent on the cellular context (e.g., protein milieu in the cell) vs. ones that are intrinsic to the DNA sequence. Here, we demonstrate that predictive models trained in one cellular context can be used to predict the MPRA output in another with reduced prediction power and that, as expected, regions whose activity is cell type specific are harder to predict in in this cross-dataset setting. We also observe that gene expression of TFs is overall consistent with the predictive ability of their binding instances, with highly expressed TFs being generally more predictive of MPRA activity. When comparing pairs of datasets for TFs that are predictive of MPRA activity, we notice that in some cases, TFs with cell type specific functionality are better predictors in that cell type.

In addition, we wanted to evaluate the applicability of our predictive models in studying the function of naturally occurring mutations. We therefore tested the ability of our framework to detect the effects of small variants – single nucleotide variants (SNV) or short insertions or deletions (indels) – on MPRA activity, and achieved similar accuracy to the state of the art methods (Zeng, et al., 2017). Finally, we applied our approach to the Regulation Saturation challenge of the Fifth Critical Assessment of Genome Interpretation (CAGI5), and demonstrate that it achieves top performance in identifying functional effects of SNVs in supervised settings.

## RESULTS

We used five publicly available MPRA datasets and one unpublished dataset collected at several labs using a range of experimental methodologies and cell types (Methods). In all cases, the MPRA constructs were designed to test endogenous human DNA sequences, and not in-silico designed synthetic sequences (Smith, et al., 2013). Thus, each element tested in each dataset is associated with a source genomic region. Each dataset consists of approximately 2,000 sequences with length that varies between 121 and 171 base pairs (Methods). Unless otherwise noted, the MPRA experiment was performed in an episomal context. The first dataset (Kwasnieski, et al., 2014), which we refer to as *K562*, consists of putative regulatory regions selected from ENCODE-based annotated regions in *K562* cells (Consortium, 2012; Ernst and Kellis, 2010; Hoffman, et al., 2013). The second and third datasets, which we refer to as *LCL-eQTL* and *HepG2-eQTL* (Tewhey, et al., 2016), consist of sequences that contain an eQTL in Lymphoblastoid Cell Lines (LCLs). The same sequences were tested in LCL and HepG2 cells, thus forming the two datasets. Notably, the *LCL-eQTL* dataset was used as the primary source for the CAGI4 eQTL causal challenge (Kreimer, et al., 2017). The fourth and fifth datasets (Inoue, et al., 2017) include candidate liver enhancers, tested in either episomal or chromosomal context. We refer to these datasets as *HepG2-epi* (for MPRA plasmids) and *HepG2-chr* (for MPRA integrated in the genome). The sixth dataset includes putative enhancer regions (F., et al., 2018), tested in chromosomal context in human embryonic stem cells (hESC). We refer to this dataset as *hESC*. Separately, for each dataset, we applied MPRAnalyze (Note S1; (Ashuach, et al.); **Methods; Figures S1-S3**), a new tool for statistical analysis of MPRA data developed in our group, to obtain (1) MPRA output: a quantitative measure of enhancer-induced transcription, computed as the ratio between the estimated abundances of transcribed RNA and the construct’s DNA. These values are estimated by constructing a nested pair of generalized linear models that extract the ratio RNA / DNA as a measure of activity while controlling for various confounding factors, and (2) a binary label that identifies active/inactive enhancers, namely enhancers whose activity significantly deviates from that of the negative controls (median-based z-score; FDR < 0.05).

### Predictive features for MPRA activity are consistent across datasets

We first defined a set of features that characterize each MPRA sequence and inspected each feature individually (**Methods**; see **Table S1** for a complete description of all features). Overall, we examined 56 features that can be divided into four categories (similarly to (Kreimer, et al., 2017)): **(1) *Experimentally measured epigenetic properties***. To define these, we mapped each assayed region to its corresponding position in the reference human genome, and then queried this position against tracks of epigenetic properties from ENCODE (Consortium, 2012). These properties were measured in multiple cell lines and include the overall number of observed TF binding sites (TFBS), histone marks, binding by chromatin structure-associated proteins (e.g., P300), chromatin accessibility (primarily by identifying DNase-hypersensitivity sites; henceforth abbreviated as DHS), and DNA-methylation. For all these features we either aggregate over all available cell types, or restrict the analysis to the same cell type in which the MPRA was conducted. **(2) *Predicted epigenetic properties***. This set of features covers similar properties as the experimentally-derived ones (e.g., TFBS or histone marks). However, instead of being directly measured, the properties are inferred based on the DNA sequence of the respective MPRA construct, using models trained on experimental data (e.g., protein binding microarrays for TFBS (Newburger and Bulyk, 2009) or ChIP-seq for histone marks (Consortium, 2012)). We use three models for this purpose: scoring of protein-DNA binding motifs (Grant, et al., 2011) and the more recent supervised methods *DeepBind* (Alipanahi, et al., 2015) and *DeepSea* (Zhou and Troyanskaya, 2015). Another feature included here is *Motif Density* – defined as the maximum number of protein-DNA binding motifs within a 20 bp window in the MPRA sequence. **(3) DNA *k-mer frequencies*** using k=5. And **(4) *Additional locus specific features***. Here we used the number of G/C in the sequence (*#GC*) as well as the length of longest polyA/T subsequence (*#polyA/T*). We also used DNA shape features (Zhou, et al., 2013) quantifying minor groove width, roll, propeller twist, and helix twist (*MGW, Roll, ProT, and HelT* respectively). Additional features in this category include: *Conservation* – evolutionary conservation score of region as predicted by phastCons (Siepel, et al., 2005). *Closest Gene Expression* – expression (TPM) of the closest gene from RNA-seq data in the corresponding cell type. *Promoter, Exon, Intron, Distal* – binary features indicating the respective location in the endogenous genome.

We use these 56 features (**Methods; Table S1**) individually in two ways: **1)** we test how well each feature correlates with the quantitative MPRA output of each dataset using seven *regression tests* (**Methods**) and **2**) we test how well each feature discriminates between active and inactive regions using two *classification tests* (**Methods**). We rank each feature for each of the nine tests and then take the median of these ranks to obtain a dataset-specific feature ranking. We then take the median across all dataset-specific ranking to obtain a global ranking of the features and sort them according to their global rank (**Figure 1**). Notably, the different statistical test are largely consistent with the global rank (**Figure 1; Table S1**), supporting its robustness. This global rank highlights chromatin accessibility (*DNase Mean*) and the number of TF binding sites (*TFBS Mean*) as the most predictive features for MPRA activity across all data sets. To gauge the robustness of our results, we repeated the above feature correlation experiments 100 times, each time sampling 80% of the loci in the data, and report the mean and standard deviation (STD) of the resulting accuracy (**Table S1**).

**Figure 1:**
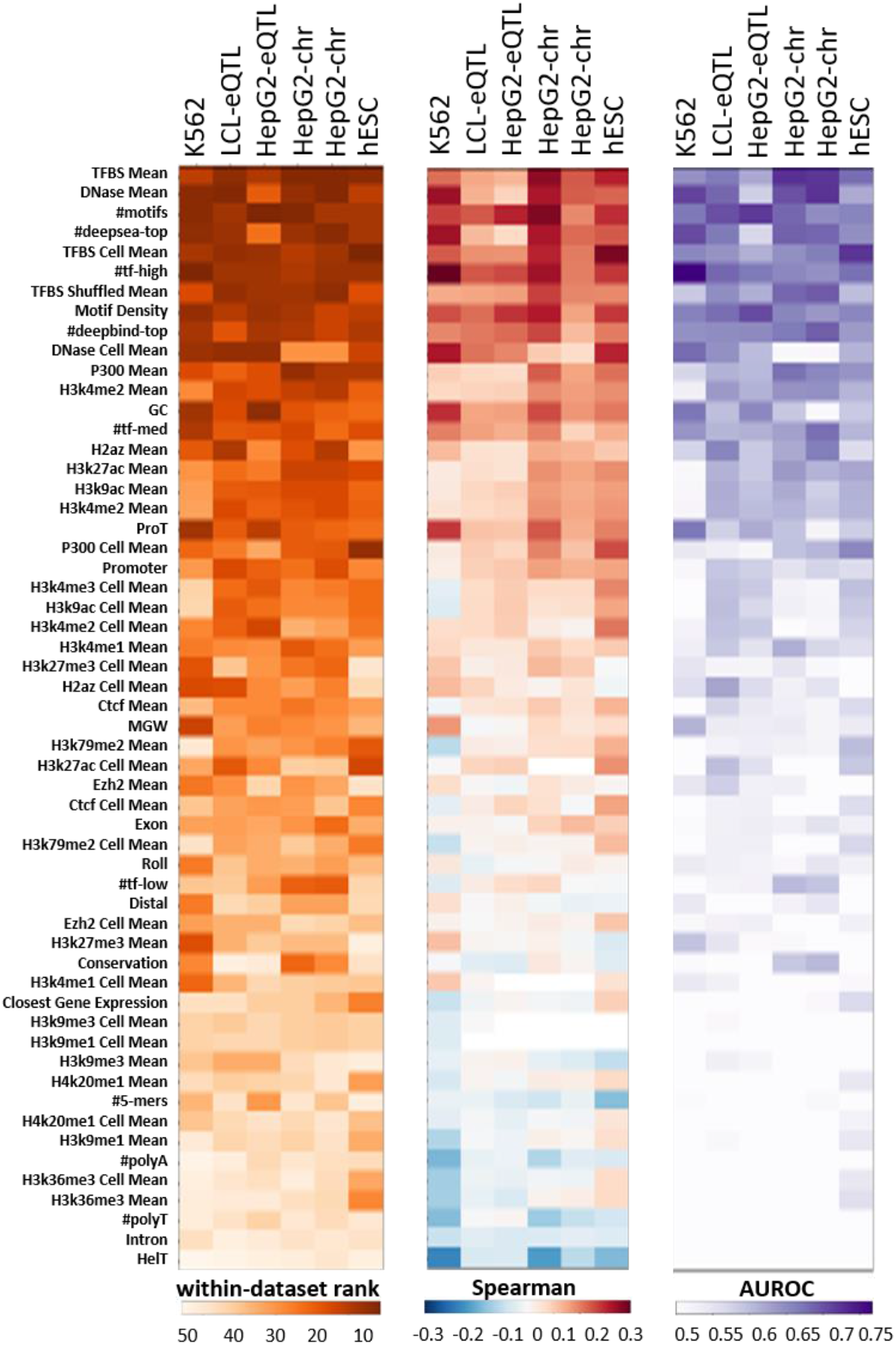
Individual feature correlation with MPRA output. The within-dataset ranking is calculated by first ranking each feature by each test, then taking the median of the regression and classification test rankings. The comprehensive ranking is the median across all the dataset rankings. The heatmaps are ordered according to the comprehensive ranking and colored according to 1) the within-dataset rank 2) the Spearman correlation coefficient for regression task 3) the AUROC value for classification task.

To further explore cell-type specificity in the context of TF binding, we stratified the TFs into three groups according to their expression level in the cell type of interest (low / intermediate / high) and sum over the number of binding sites in each group. While these three features *#tf-high, #tf-med, #tf-low* had a strong correlation (especially *#tf-high)* with MPRA activity (**Figure 1**), they are still less predictive than *TFBS Mean* (the simple mean across all TFBS-related features). Consistently, we found several cell-type agnostic features such as *GC* content and *#motifs* that are predictive of MPRA activity as well (**Figure 1; Figures S4-S5**).

Furthermore, we found that limiting the set of TFs in a manner specific to the cell-type under investigation (e.g. for the *K562* dataset, *TFBS Cell Mean* only considers TF ChIP-seq experiments conducted in K562 cells) does not improve accuracy (**Table S1**), compared with taking all available data regardless of cell type of origin (*TFBS Mean*). This observation is consistent with previous work on enhancer annotation, showing that integration of diverse datasets from different cellular contexts improves developmental enhancer prediction over approaches based on single context data (Erwin, et al., 2014). As additional control, we randomly subsampled N (the number of TFs used to calculate *TFBS Cell Mean*) ChIP-seq experiments that were conducted in a cell type different from the one used for MPRA, and computed the mean number of binding sites. Consistent with the results above, we found that the predictive capacity of this random set of TFs binding scores (considering 100 randomly selected sets for each of our six data sets; denoted *TFBS Shuffled Mean)* is not lower than that of ChIP-seq experiments conducted in cell type in which MPRA was conducted (empirical p-value >0.25).

### Predictive models of MPRA activity are similar across datasets

We next turned to the construction of supervised models that are trained to predict the MPRA output either as a quantitative measure of enhancer activity (i.e. regression task) or as a binary label that distinguishes between active and inactive enhancers (i.e. classification task). To this end, we considered a collection of regression models (*Elastic Net* (Hui Zou, 2005), *Random Forest* (Breiman, 2001), *Extra Trees* (Geurts, et al., 2006), *Gradient Boosting* (Zhu, et al., 2009), and *ensemble*) and classification models (*Random Forest* (Breiman, 2001), *Extra Trees* (Geurts, et al., 2006), *ensemble*), which we applied separately for each data set. We trained these models using a set of features that extends the one investigated in **Figure 1**, with the following categories: **(1) *Experimentally measured epigenetic properties*** – 1095 binary features based on ENCODE data (Consortium, 2012). These features indicate whether the genomic region overlaps with experimentally measured tracks of: TFBS from ChIP-seq experiments, histone modifications, and DNase-hypersensitivity sites across different cell types (**Table S1**). **(2) *Predicted epigenetic properties***. This set consists of three sources: *(i) DeepBind* – 515 features, each indicating a binding score of a certain TF, predicted by a sequence-based neural network model trained on protein-binding microarrays (Alipanahi, et al., 2015). (ii) *DeepSea* – 919 binary features, indicating predictions of various events related to chromatin structure, namely TF binding, DNA accessibility, and histone modifications. These events were predicted by a sequence-based neural network model trained on ENCODE data (Zhou and Troyanskaya, 2015). (iii) *Motifs* – 2065 binary features indicating motif hits (Consortium, 2012; Kheradpour and Kellis, 2014) (Grant, et al., 2011). **(3) DNA *k-mer frequencies*** – 1024 binary features, indicating the presence or absence of all possible nucleotide 5-mers. **(4) *Additional locus specific features*** as in Figure 1 (**Table S1**).

We evaluate the accuracy of prediction in each combination of data set x prediction method x feature category using 10-fold cross validation. We report the mean and standard deviation (STD) of the resulting scores (**Figure 2**). Importantly, we do not use our evaluation of individual features in Figure 1 during model training (e.g., for feature selection), thus avoiding circularity.

**Figure 2:**
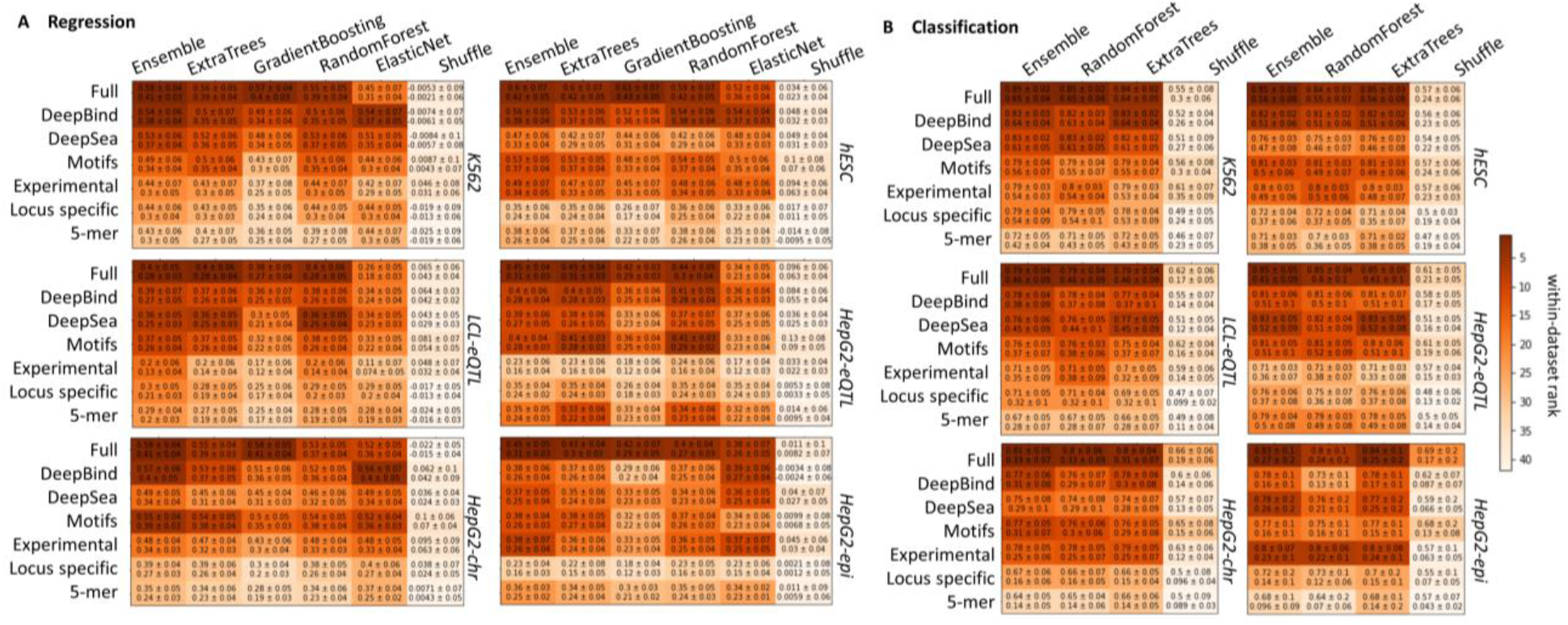
Performance of (A) regression models and (B) classification models with different feature combinations. The within-dataset ranking is calculated for each cell by taking the median of the rankings for all the (A) regression or (B) classification tests within a dataset. Each heatmap is colored according to the within-dataset rankings. The statistics are *mean ± std* for (A) Spearman and Kendall tests or (B) AUROC and AUPRC tests.

Reassuringly, the accuracies of our top model for predicting MPRA activity on the *LCL-eQTL* dataset (regression and classification: 0.4 Spearman correlation and 0.79 AUROC respectively) matched that of the top ranking group in the CAGI4 challenge (0.34 Spearman correlation and 0.8 AUROC) (Zeng, et al., 2017). Consistent with our results for single features, we observe an overall agreement in our results across datasets, both in terms of the relative performance of each algorithm, and in terms of the importance of each feature category. Specifically, we observe that non-linear methods perform better (e.g., compare elastic net to random forest) and that an ensemble approach (aggregating over all classifiers or regression methods) tends to have the highest performance (**Figure 2; Table S2**). Among the feature categories, the predicted TF binding properties according to *DeepBind* are top performers, and the union of all feature categories generally yields the best performance, indicating that even with a large feature set the various models still do not over-fit. To further test this, we trained our models on shuffled labels (**Methods**), and observed that the performance significantly decreases in all cases, including the more complex ensemble model that uses the complete feature set.

Another result consistent with the ones observed with single features regards the importance of cell type specificity, where we again noticed that limiting the epigenetic features to be cell type specific does not increase accuracy (**Figure S6; Table S3**). Finally, it is interesting to note that the accuracy achieved with a chromosomal MPRA library in HepG2 cells (*HepG2-chr*) tends to be slightly higher than the one obtained with an episomal library (*HepG2-epi*) (regression: 0.59 vs. 0.45 Spearman correlation and 0.41 vs. 0.31 Kendal correlation; **Figure 2**). These results are consistent with a recent comparison between these two experimental approaches (Inoue, et al., 2017) that found chromosomal MPRA to be more reproducible, have higher correlation with epigenetic marks and work in variety of cell-types that are harder to transfect (e.g. hESCs); however, more datasets are required to substantiate this finding.

### Transferring knowledge between cell types

Using existing MPRA data to build models that can be applied across different cellular backgrounds and for genome-wide predictions of regulatory elements can be useful for prioritizing functional regulatory regions, which can guide the design of new MPRA panels and used for analysis purposes. To evaluate how well our models generalize to a new cellular context where MPRA data is not available, we tested the extent to which models trained in each dataset can be used to predict the outcome in the remaining datasets. Based on the results in Figure 2, we take the *Full* set of features (i.e., all feature categories) and use the *ensemble* model for both the regression and classification tasks. We avoid training on any genomic region from one dataset (e.g. *LCL-eQTL*) that is already in the test set from another dataset (e.g. *HepG2-eQTL*).

We observe that the accuracy of prediction is reduced in this cross-dataset setting, compared with the cross validation setting (e.g. cross validation for K562: 0.58 Spearman; **Figure 2**; comparing to cross-dataset learning: 0.23, 0.21, 0.44, 0.3, 0.33 Spearman for *LCL-eQTL, HepG2-eQTL, HepG2-chr, HepG2-epi, hESC* respectively; **Figure 3; Table S4**). However, the performance is generally robust for determining if a region is active (e.g. cross validation for K562: 0.85 AUROC; **Figure 2**; comparing to cross-dataset learning: 0.7, 0.67, 0.75, 0.74, 0.68 AUROC for *LCL-eQTL, HepG2-eQTL, HepG2-chr, HepG2-epi, hESC* respectively; **Figure 3; Table S4**). These results suggest that MPRA data in one cellular context can be leveraged to distinguish between regions of regulatory importance in another.

**Figure 3:**
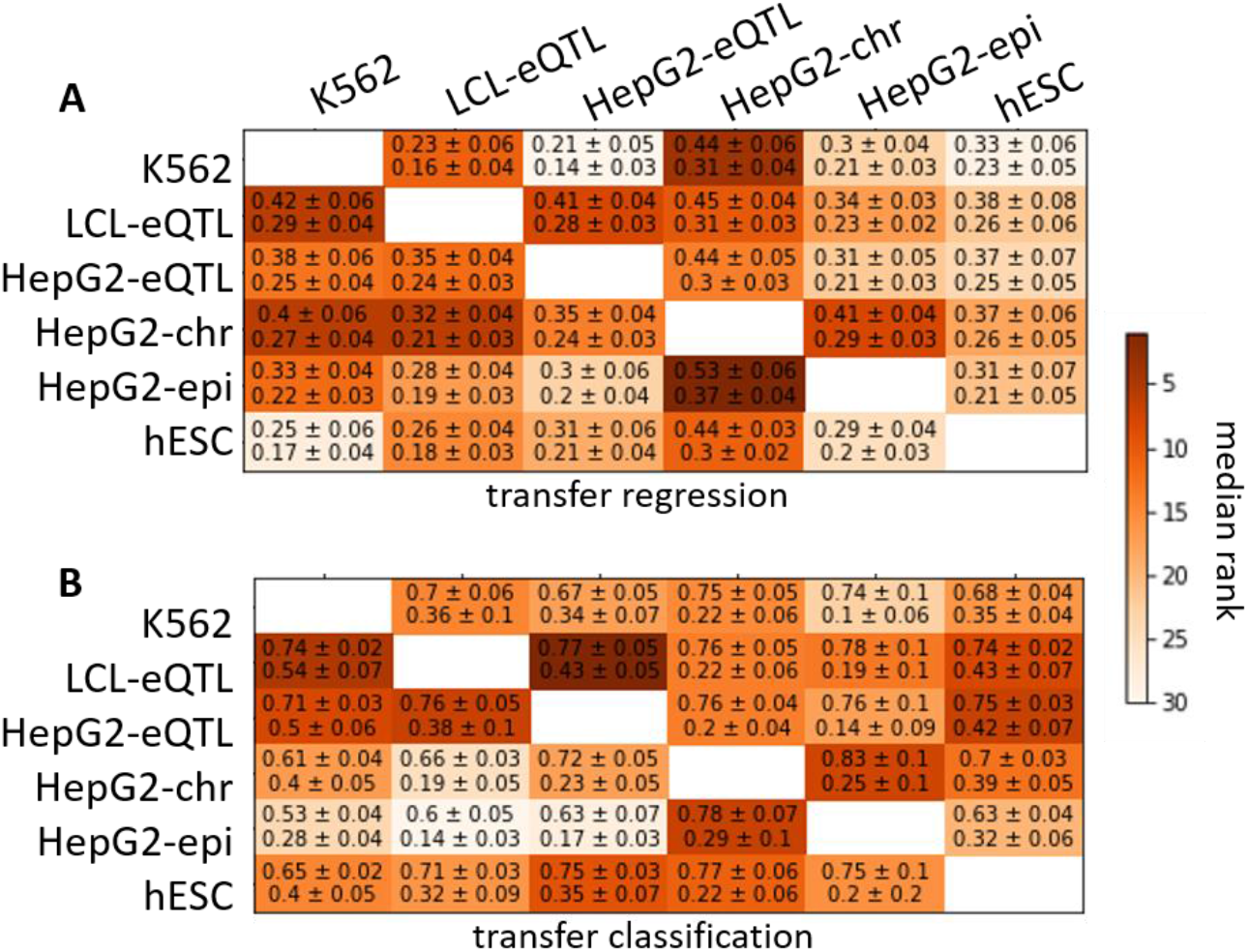
Performance of cross-dataset learning for (A) regression task and (B) classification task between cell types. All cross-dataset learning models are ensemble models: with full features. Each cell is colored according to the median over the ranks of all (A) regression tests or (B) classification tests. The statistics are *mean ± std* of (A) Spearman and Kendall tests or (B) AUROC and AUPRC tests.

We hypothesized that genomic regions that are uniquely active in a certain cell type would be harder to predict in a cross data set setting. To explore this, we took advantage of the *LCL-eQTL* and *HepG2-eQTL* datasets, which include the same set of genomic regions. We first examined the distribution of three region categories in these two datasets (Figure 4A): common regions (i.e. active regions in both datasets), cell type specific regions (i.e. regions active in one of the datasets), and inactive regions (i.e. regions not active in both datasets). We then examined prediction performance for each of the region categories (Figure 4B) in cross-data set analysis where we apply the classifier built on one dataset to annotate regions in the other dataset as active or not. To assess this, we defined the “hardness” of the region based on the difference between the predicted score (in range [0, 1]) and the class label (1 for active and 0 for not-active region). Reassuringly, we observe that cell type specific regions are harder to predict in cross-dataset learning (Figure 4B). These results suggest that while the MPRA signal can be predicted to some extent using cell type agnostic components, it also depends on cell type specific ones. Interestingly, and consistent with our cross validation (i.e., per-dataset) analysis, we observe that the cross-dataset accuracy achieved with models trained on chromosomal MPRA library (*HepG2-chr*) is higher (0.33, 0.28, 0.3, 0.31 Spearman and 0.53, 0.6, 0.63, 0.63 AUC for K562, *LCL-eQTL, HepG2-eQTL, hESC* respectively) than the one obtained with an episomal library (*HepG2-epi*) (0.4, 0.32, 0.35, 0.37 Spearman and 0.61, 0.66, 0.72, 0.7 AUC for K562, *LCL-eQTL, HepG2-eQTL, hESC* respectively). (Methods; Figure 3; Table S4).

**Figure 4:**
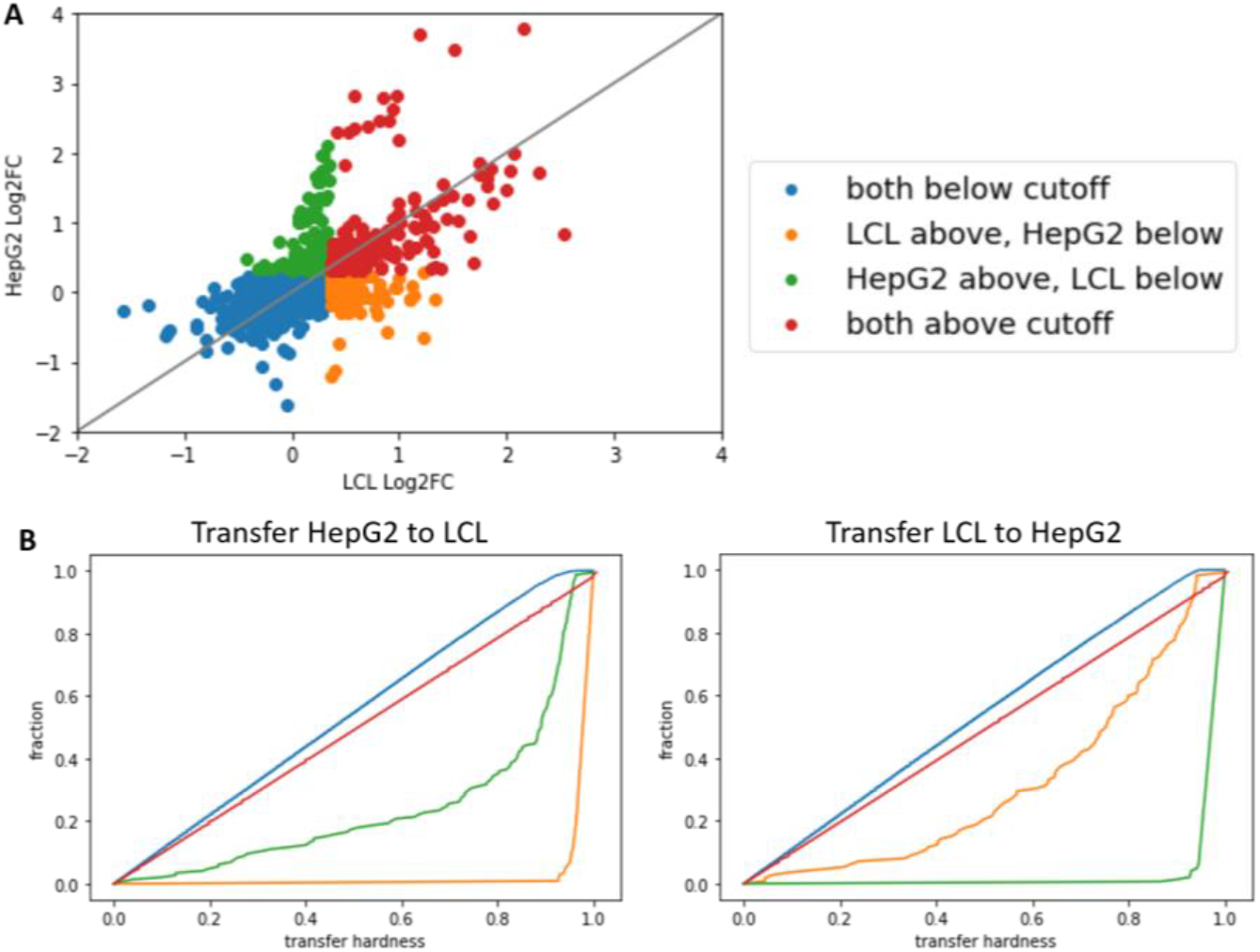
(A) *LCL-eQTL* vs. *HepG2-eQTL* MPRA activity by log2 alpha values. The points are colored according to activity in each of the datasets (active/inactive is defined as above/below 1.5 cutoff respectively). (B) We define hardness as the rank-normalized absolute difference between the ground truth binary activity label (0 or 1) and predicted probability. The cumulative distribution function of the hardness for each of the four activity groups when training the ensemble, full feature cross-dataset model on *HepG2-eQTL* (Left subfigure) and *LCL-eQTL* (Right subfigure), and testing on *LCL-eQTL* and *HepG2-eQTL*, respectively.

### Contributions of individual TFs to the accuracy of predicting MPRA outcome

We wanted to explore which factors in different cells drive the activity of regulatory regions, and hypothesized that the protein milieu in the cell might act as one. To this end, we examined the contribution of individual TFs to MPRA activity. We recorded the correlation between each TF binding signal (*DeepBind* prediction) and the activity of each MPRA region (Alipanahi, et al., 2015). Similarly to our analysis in **Figure 1**, we then ranked the TFs based on their predictive ability across datasets, thus revealing several TFs whose binding is generally informative of regulatory activity of MPRA constructs in all cellular contexts in this study (**Figure 5; Figure S7-8; Table S5**). For instance, two TF families with a dataset-wide high predictive capacity, that is also supported by experimentally-evaluated binding from ChIP-seq (**Figure S8; Table S5**) sites are JUN and FOS. Proteins of the FOS family dimerize with proteins of the JUN family, thereby forming the transcription factor complex AP-1, which has been implicated in a wide range of cellular processes, including cell growth, differentiation, and apoptosis across different cell types (Ameyar, et al., 2003). More generally, we find that TFs whose binding is commonly predictive of MPRA activity across data sets are also highly expressed across all the three cell types, as indicated by RNA-seq data (**Figure 5**). Indeed, the gene expression of TFs is overall consistent with their predictive capacity, whereby more predictive factors have overall higher expression as measured by RNA-seq (Consortium, 2012) (**Figure 5** – right four columns) across all cell types (Wilcoxon rank sum test of top vs. bottom 50 factors: p-value of 3.9e-6, 1.36e-5, 8.7e-4, and 8.0e-4 for K562, LCL, HepG2, and H1hESC respectively).

**Figure 5:**
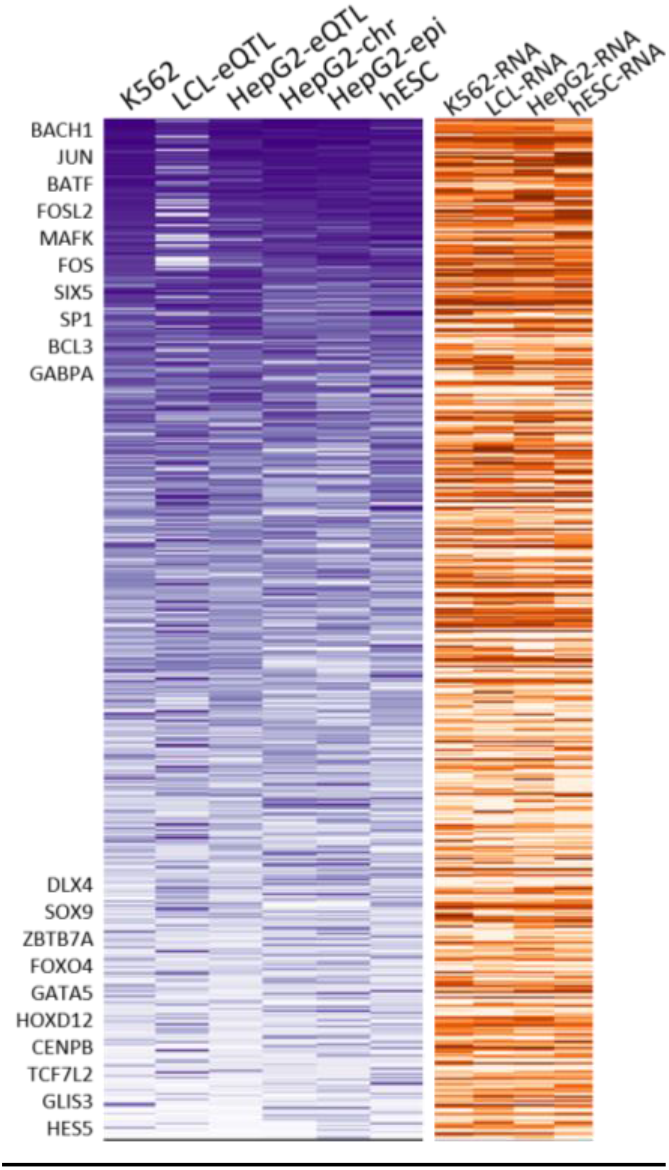
Contribution of individual DeepBind TF binding for predicting regulatory activity of MPRA constructs. The within-dataset ranking is calculated by taking the per feature median rank across all classification and regression tests. The comprehensive ranking is the per feature median overall within-dataset rankings. TFs are sorted from best (smallest) to worst comprehensive rank. (Left) Heatmap of the within-dataset rankings. (Right) The per TF ranking of its mRNA levels measured by RNA-seq in each of the four cell lines. Names of the common top/bottom 10 factors are indicated on the left.

### Exploring common and distinct TF binding between datasets

We next proceeded to explore TFs whose binding is predictive of MPRA activity only in specific cell types. To this end, we defined, for each dataset, a set of *predictive TFs*, as the set of bound TFs (predicted by *DeepBind*) that is significantly correlated with MPRA output (Spearman FDR corrected p-value < 0.05). We then compare across pairs of datasets to determine if there is significant overlap in *predictive TFs*. To this end, for each pair of datasets, we calculated the fold enrichment of the overlap between the predictive TF set, and evaluated the significance of this overlap using a hypergeometric p- value (**Figure 6A**). Overall, we see that there is significant overlap across every pair of datasets. Interestingly, the similarity between datasets seem to be dominated by the similarity between the MPRA sequences and less so by the similarity in cellular context. Specifically, the *HepG2-chr* and *HepG2-epi* pair and *LCL-eQTL* and *HepG2-eQTL* pair had the strongest overlap, suggesting that the same genomic regions tested in different conditions have correlated signals in MPRA. However, this result may depend on the specific sequences studied, and further data needs to be collected to substantiate it.

**Figure 6:**
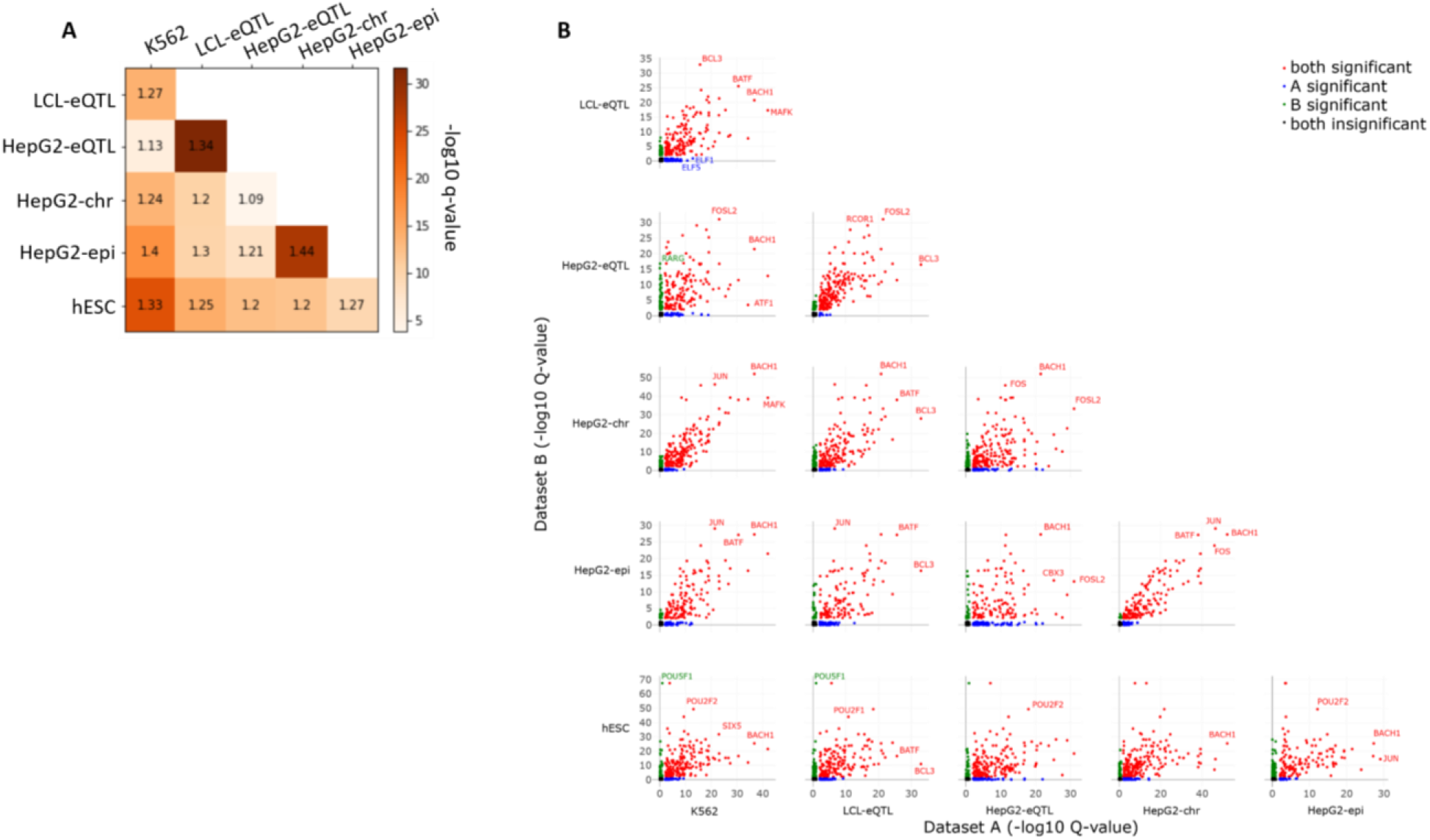
Similarities and differences in TFs whose binding is predictive of MPRA activity between datasets. For each dataset, we define a set of *predictive TFs*, as the set of bound TFs (predicted by *DeepBind*) that is significantly correlated with MPRA output (Spearman q-value< 0.05). (A) Similarity of *predictive TFs* between datasets. For each pair of datasets, a hypergeometric test is performed on the sets of *predictive TFs* of both datasets, resulting in a q-value indicating the likelihood of the overlap occurring by chance (color scale). We also calculate the enrichment ratios of *predictive TFs* for each pair of datasets (cell text). (B) Differences in *predictive TFs* between datasets. For each dataset, we only plot high confidence *predictive TFs* (i.e. significant TFs) that have Spearman q-values of less than 0.01, and *non-predictive TFs* (i.e. non-significant TFs) have q-values of greater than 0.1.

We further examine the *predictive TFs* that differ between pairs of datasets (**Figure 6B; Table S6**), and provide a list of top *predictive TFs* in at least one dataset. In some cases, we find proteins whose function is related to the cell type under investigation. For instance, when comparing the two datasets with the lowest *similarity score* for *predictive TFs, K562* to *HepG2-eQTL*, we find that RARG (a retinoic acid receptor which belongs to the nuclear hormone receptor family and is associated with liver risk phenotype (Roberts, et al., 2010)) is predictive in *HepG2-eQTL* but not *K562*. When comparing *K562* to *LCL-eQTL*, we observed that the genes in the ETS family (ELF1, ELF5, ELF3, ETV6, ELK3) are predictive only in *K562*. These genes are known to be expressed in hematopoietic tissues and cell lines, and play a role in hematopoietic cell development (Clausen, et al., 1997). When comparing *hESC* to the other datasets, we observe a known pluripotent factor-POU5F1 (Boyer, et al., 2005) to be predictive only in *hESC* for most of the comparisons (**Table S6**).

Overall, these results support the notion that both sequence-intrinsic and cell-type specific properties are determining MPRA activity. We also find that the cell-type specific component may be captured by the activity of TFs whose function is associated with the cell-type under investigation.

### Studying the effects of small genetic variants on MPRA output with application to CAGI challenges

MPRA can be used to study the transcriptional effects of small variants that commonly occur in regulatory regions, namely SNPs and small indels (Tewhey, et al., 2016). We wanted to examine if we can predict these effects in the synthetic setting of MPRA. An important feature of the *LCL-eQTL* and *HepG2-eQTL* datasets (Tewhey, et al., 2016) is that each of the sequences (which come from the reference human genome) is matched with an alternative allele (single nucleotide variants (SNVs) or short indels) (Lappalainen, et al., 2013) that was tested by MPRA as well. Here, we test the ability of our models to determine the amount of shift in MPRA transcriptional activity, comparing each reference allele to its alternative. We focus on the *LCL-eQTL* dataset, which was featured in the CAGI4 challenge, and for which the results of competing methods are available (Kreimer, et al., 2017). To this end, we first applied the ensemble regression model above to predict transcriptional activity of the reference and alternative alleles, separately. Next, we trained a logistic regression using the absolute difference between those predicted expression values as a feature to predict whether there is a significant allelic variation. This strategy leads to favorable results (0.67 AUROC, 0.45 AUPRC in 5-fold cross validation), compared to other participants in the CAGI4 challenge (best result: 0.65 AUROC, 0.45 AUPRC). Unsurprisingly, however, the absolute performance is substantially lower than that achieved in the task of predicting the transcription of individual sequences, which can be expected as this task relates to a much more nuanced signal.

Importantly, our approach is also one of the top performing methods in the CAGI5 Regulation Saturation Challenge, also titled “Predicting individual non-coding variant effects in disease associated promoter and enhancer elements.” This challenge experimentally assessed the effects of 17,500 SNVs in 5 enhancers (IRF4, IRF6, MYC, SORT1, ZFAND3) and 9 promoters (F9, GP1BB, HBB, HBG, HNF4A, LDLR, MSMB, PKLR, TERT) of lengths 187 to 600 bps with saturation mutagenesis MPRA. Participants were asked to predict the directional effect (negative, zero, positive) of a SNV on the MPRA signal (in this case - log ratio of RNA counts to DNA counts).

We featurized each variant and wildtype sequence with the subset of *Full* features that differ between variant and wildtype. We adjusted our *ensemble* approach to concatenate the sets of features from the variant and wild-type sequences to predict the directional effect via multiclass classification (**Methods**). For the “direction” prediction, our method yielded the best correlations (0.318 Pearson and 0.249 Spearman), as well as competitive AUROCs (0.762 for positive vs negative, 0.706 for positive vs. rest, 0.776 for negative vs rest) (**Table S7**).

## DISCUSSION

MPRA holds a great promise to be a key functional tool that will increase our understanding of gene regulatory elements and the consequences of nucleotide changes on their activity. While previous studies already used MPRA to construct predictive models of transcriptional regulation, its generalizability across cellular contexts and its applicability for studying the endogenous genome have not yet been systematically evaluated. Here, we study MPRA data from a number of cellular systems to determine which features are reflective of the cellular context (e.g., protein milieu in the cell), and which are intrinsic to DNA sequence. We aimed to incorporate the most recently produced MPRA datasets of endogenous sequences in this work, but had to exclude several datasets after quality control analysis (e.g. The data in (Maricque, et al., 2017) consisted of few barcodes per candidate enhancer and had significant inconsistency across replicates. The experimental design in (Ulirsch, et al., 2016) included three genomic regions per enhancer, in overlapping windows. Activity measurements were highly variable between windows of the same enhancer, while many of the features we use were shared among the overlapping windows). We explore the extent by which knowledge on regulatory activity in one cellular context can be used to make predictions in a held out cellular context. Finally, we examine the ability of our framework to detect the effects of small variants on MPRA activity. Our results represent, to the best of our knowledge, the first such comprehensive analysis.

Our work highlights genome accessibility and TF binding as the strongest predictors of regulatory activity, with no observed advantage to cell type specific features. When applying prediction models, we observe that performance is improved when using an ensemble of all features, with no significant prediction improvement when using cell type specific features. These results imply that part of the signal observed in MPRA studies is not cell type specific. Interestingly, models trained with chromosomal MPRA data yield better predictions across datasets than those trained on episomal MPRA data, stressing the importance of this experimental approach that conveys a more reliable representation of the endogenous settings.

When training on one cell type and predicting on another cell type, we observe overall lower but robust results, with regions enriched in cell type specific signal being harder to predict. Notably, we detect a communal component across datasets with a group of TFs being top predictors, as well as some cell-specific factors that seem to be involved in phenotypes associated with the corresponding cell type. In the MPRA setting the *cis* environment (e.g. chromatin) is altered, thus generally not cell-type specific, and the *trans* environment (e.g. TF binding) remains similar, hence we can still observe predictive factors that are cell type specific.

As seen through its performance in the CAGI5 Regulation Saturation challenge, our approach is competitive in the high resolution task of predicting the functional effects of SNVs in a supervised setting.

Our work provides a comprehensive resource of annotation for thousands of endogenous sequences across the genome. Furthermore, we demonstrate the performance of different machine learning models for MPRA activity prediction (**Tables S2-3**) by using publicly available tools. Our approach can highlight functionally important regulatory regions across the genome in a cell-type agnostic fashion and can be leveraged for an efficient design of future MPRA experiments by prioritizing regions of interest.

## ACKNOWLEDGMENTS

This work was supported by the National Institute of Mental Health (NIMH) grant number 1K99MH117393 (A.K). A.K, Z.Y. and N.Y. were supported by the National Human Genome Research Institute (NHGRI) grant number U01HG007910. N.A. is supported in part by the NHGRI grant number 1UM1HG009408. The CAGI experiment coordination is supported by NIH U41 HG007346 and the CAGI conference by NIH R13 HG006650.

## METHODS

### MPRA datasets

We used five publicly available MPRA datasets and one unpublished dataset. (1) ***K562*** – putative regulatory regions (Kwasnieski, et al., 2014) selected from ENCODE-based annotated regions in K562 cells (Consortium, 2012; Ernst and Kellis, 2010; Hoffman, et al., 2013). This set includes 600 regions annotated as enhancers, 600 as weak enhancers, 300 as repressed in K562 cell line, 600 enhancer predictions from the H1hESC cell line that are not annotated as weak enhancers or enhancers in K562 cells, and 1,136 negative controls – random sequences from each class above were chosen and scrambled while maintaining dinucleotide content. The regions range from 121 to 130 base pairs and were tested in episomal context in K562 cells. Data from all sequences were used to fit MPRAnalyze, although only the 1,500 regions annotated with the K562 cell-line were used in the remaining analyses. (2) ***LCL*-eQTL** – 78,738 regions (Tewhey, et al., 2016) that contain an eQTL in Lymphoblastoid Cell Lines (LCLs), 150 base pairs, tested in episomal context in LCL. (3) ***HepG2-eQTL*** – the same set of elements (Tewhey, et al., 2016) as above, tested in episomal context in HepG2 cell line instead of LCL. For both datasets 2 and 3, all of the 78,738 regions were used to fit MPRAnalyze, while 3,044 regions corresponding to the first test group in the CAGI4 challenge (Kreimer, et al., 2017) were used for the remaining analyses. (4) ***HepG2-chr*** – 2,236 candidate liver enhancers (Inoue, et al., 2017) and 102 positive and 102 negative control sequences. Each sequence is 171 base pairs and tested in chromosomal context. (5) ***HepG2-epi*** – the same set of elements (Inoue, et al., 2017) as above, tested in episomal context. For both datasets 4 and 5, all regions were used to fit MPRAnalyze and the 2,236 candidate enhancer regions were used for the remaining analyses. (6) ***hESC*** – 2,464 putative enhancer regions (F., et al., 2018) and 200 negative controls. Each region is 171 base pairs and tested in chromosomal context in hESC cell line. All regions were used to fit MPRAnalyze, while only the 2,268 candidate enhancer regions were used for the remaining analyses.

### Quantifying activity of regions using MPRAnalyze

For each dataset, we obtain the RNA and DNA raw counts for each barcode. We obtain a quantitative measure of enhancer-induced transcription using MPRAnalyze ((Ashuach, et al.); **Note S1**). MPRAnalyze assumes a linear relationship between the RNA and DNA counts, with the scaling parameter, denoted *alpha*, as the transcription rate. The method uses a parametric graphical model to incorporate external covariates and dispersion estimates into quantifying alpha.

The MPRAnalyze model assumes the DNA counts are Gamma-distributed and that given the latent plasmid count, the RNA counts are Poisson-distributed centered around the product of the plasmid count and alpha. This results in a closed-form negative-binomial likelihood function for the RNA counts. External covariates such as barcode effect, batch effects and conditions of interest are then incorporated into the model by constructing a pair of nested generalized linear models: one using the DNA counts to estimate the latent plasmid counts, and the other using these latent plasmid counts along with the RNA raw counts to estimate alpha.

Classification of active / inactive enhancers is done by using the fitted alpha values. If a dataset has control regions (*K562* and *hESC*), we first calculate a robust version of the standard score from the alpha values by subtracting the median over the control regions and dividing by the median absolute deviation (MAD) of the control regions. If no control region exists for the dataset, we perform the previous step with the median and MAD over all regions instead of just the control regions. We then compute the survival function for each standard score and apply the Benjamini-Hochberg (BH) correction. The active regions are then defined as regions with a false discovery rate (FDR) of less than 0.05.

### Features

We assessed the correlation of 56 single features (**Table S1**) with MPRA activity.

(a) ***#GC; #polyA, #polyT*** – number of G/C in the sequence; length of longest polyA/T subsequence. (b) ***#5-mers*** – number of distinct 5-mers in the sequence. (c) ***MGW, Roll, ProT, HelT*** – DNA shape features (Zhou, et al., 2013) quantifying minor groove width, roll, propeller twist, and helix twist. (d) ***Conservation*** – evolutionary conservation score of region as predicted by phastCons (Siepel, et al., 2005). (e) ***Closest Gene Expression*** – expression (TPM) of the closest gene from RNA-seq data in the corresponding cell type. (f) ***Promoter, Exon, Intron, Distal*** – binary features indicating whether the element intersects a promoter, exon, and intron. *Distal* is defined to be 1 if the element does not intersect with either promoter, exon or intron annotations. (g) ***#motifs, Motif Density*** – number of significant DNA-binding ENCODE motifs (Consortium, 2012) from simple DNA-binding motif scoring (Grant, et al., 2011), maximum number of motifs within a 20 bp window in the sequence. (h) ***#deepsea-top, #deepbind-top*** – number of TFs quantifications above 90^th^ percentile across all the regions predicted by *DeepSea* / *DeepBind*. (i) ***#tf-high, #tf-med, #tf-low*** – number of TFs that are bound above 90^th^ percentile by *DeepBind* and rank in the top, middle, or bottom 100 (out of 515) for RNA-seq TPM in the relevant cell type. (j) ***<factor> [Cell] Mean, TFBS Shuffled Mean*** – mean across subsets of *Experimental* features. <factor> can be *TFBS, DNase, Ctcf, Ezh2, H2az, H3k4me1, H3k4me2, H3k4me3, H3k9ac, H3k9me1, H3k9me3, H3k27ac3, H3k27me3, H3k36me3, H3k79me2, H4k20me1, P300*. For these factors we take the mean of the binary overlaps over all corresponding [, cell-type specific to the dataset’s cell-type,] *Experimental* features. *TFBS Shuffled Mean* is the mean across *n* **non** cell-type specific, randomly chosen *TFBS* features, where *n* is the number of features in *TFBS Cell Mean*.

### Statistical tests

We examine the predictivity of features and accuracy of prediction models using several statistical tests. For regression task – e.g. predicting quantitative activity – we applied several correlation measures (Pearson, Spearman, Kendall) considering either the entire test data or regions at the top 25% of quantitative activity; we also applied another Spearman correlation test after first binning quantitative activity by quintiles. We refer to these seven tests as the *regression tests*. For classification task – e.g. predicting active or not active – we record the AUROC (area under receiver operating characteristic curve) and AUPRC (area under precision recall curve); we refer to these two tests as the *classification tests*. The significance of each *regression task* was evaluated by the respective statistical test *q-values*, which are obtained from *p-values* via the Benjamini–Hochberg correction. The significance of classification was evaluated by the *q-values* of the Kolmogorov-Smirnov test on the predictions with positive ground truth labels.

### Training and testing

We deterministically divide each dataset into 10 sections; datasets with the same regions (*LCL-eQTL* and *HepG2-eQTL*, and *HepG2-chr* and *HepG2-epi*) are divided consistently. For the supervised case, we perform 10-fold cross-validation where each fold trains the model on 9 training sections then evaluating on the remaining section. For the cross-dataset case, we perform 10-fold cross-validation where each fold trains the model on 9 sections from the training dataset, then evaluating on the corresponding remaining section in the last dataset. We use the statistics from each fold to calculate the overall mean and standard deviation statistics.

When comparing cross-dataset learning performance between training on chromosomal MPRA data (*HepG2-chr*) vs. training on episomal MPRA data (*HepG2-epi*), we observe that training on *HepG2-chr* showed better results than *HepG2-epi* 37 out of 40 times (comparing results across different statistical tests) (Figure 3; Table S4). Same regions were used for training and testing was done on the other four datasets.

### Prediction models description

We predict the quantitative activity from element features with four regression models and their ensemble. The four models are a linear regressor with ElasticNet regularization (Hui Zou, 2005) with 0.5 as the L1 and L2 regularization coefficients and a RandomForest regressor (Breiman, 2001), an ExtraTrees regressor (Geurts, et al., 2006), and a GradientBoosting regressor (Zhu, et al., 2009), each with 1000 estimators. The ensemble method is implemented by taking the average prediction of all four regression models.

For the classification task, we use a RandomForest classifier (Breiman, 2001) and an ExtraTrees classifier (Geurts, et al., 2006), each with 1000 estimators, as well as their ensemble. The ensemble method averages the predicted probability from each classifier.

For both regression and classification, we define a shuffle model with the same composition as an ensemble model but shuffles the labels of the training set before training. This allows us to quantify the probability of producing our ensemble results by chance.

### CAGI5 model description

We predicted the directional effects (positive, zero, negative) of 13,186 SNVs from 5 enhancers and 9 promoters after training on 4,650 different SNVs from the same enhancers and promoters. For each SNV, we obtain the variant and wildtype sequence, each of length 187 to 600, then featurize both variant and wildtype with the 4,535 features that differ between variant and wildtype: *Predicted epigenetic properties, DNA k-mer frequencies, #GC, #polyA/T*, DNA shape features, and *conservation*. We concatenate the features from variant and wildtype into a feature vector of size 9,070.

We split the set of SNVs into two sets: one containing all enhancer SNVs and one containing all promoter SNVs. We train a separate ensemble of 5 RandomForest classifiers and 5 ExtraTrees classifiers for each set to predict the direction class (positive, zero, negative). Each classifier consists of 1000 estimators, and each estimator considers the square root of the total number of features when looking for the best split.

